# The effects of a TMS double lesion to a cortical network

**DOI:** 10.1101/517128

**Authors:** Ian G.M. Cameron, Andreea Cretu, Femke Struik, Ivan Toni

## Abstract

Transcranial magnetic stimulation (TMS) is often used to understand the function of individual brain regions, but this ignores the fact that TMS may affect network-level rather than nodal-level processes. We examine the effects from a “double lesion” to two frontoparietal network nodes compared to the effects from single lesions to either node. We hypothesize that the absence of additive effects indicates that a single lesion is consequential to a network-level process. Twenty-three humans performed pro- (look towards) and anti- (look away) saccades after receiving continuous theta-burst stimulation (cTBS) to right frontal eye fields (FEF), dorsolateral prefrontal cortex (DLPFC) or somatosensory cortex (S1) (the control region). On a subset of trials, a TMS pulse was applied to right posterior parietal cortex (PPC). FEF, DLPFC and PPC are important frontoparietal network nodes for controlling anti-saccades. Bayesian T-tests were used to test hypotheses for additive double lesion effects on saccade behaviors (cTBS plus TMS pulse) against the null hypothesis that double lesion effects are not different than single lesion effects. We observed strong evidence (BF_10_ = 325.22) that DLPFC cTBS plus PPC TMS lesion enhanced impairments in ipsilateral anti-saccade amplitudes over DLPFC cTBS alone, but not over the effect of the PPC pulse alone (BF_10_ = 0.75). Therefore, effects were not additive, and no other evidence for additive effects was found (BF_10_ < 3). This suggests that saccade-control computations are distributed across this network, with some degree of compensation by PPC for the DLPFC lesion.

## Introduction

It is well known that the effects of transcranial magnetic stimulation (TMS) extend beyond the site of stimulation (Ilmoniemi et al., 1997; Ko et al., 2008; Morishima et al., 2009; Paus et al., 1997; Ruff et al., 2006). In some instances, distal effects may reflect compensatory responses to the TMS lesion (Hartwigsen et al., 2013; O’Shea, Johansen-Berg, Trief, Göbel, & Rushworth, 2007; Sack, Camprodon, Pascual-Leone, & Goebel, 2005), suggesting “homeostatic metaplasticity” (Müller-Dahlhaus & Ziemann, 2015) at the level of a network node. Here we assess another functionally relevant possibility: whether behavioral consequences of a spatially-localized perturbation from TMS are driven by the distributed nature of computations throughout a circuit (Price & Friston, 2002).

The saccadic eye-movement system provides a testing ground for assessing circuit-level consequences of TMS (Leigh & Kennard, 2004; Munoz, Armstrong, & Coe, 2007). Roles of three cortical nodes, frontal eye fields (FEF), dorsolateral prefrontal cortex (DLPFC), and posterior parietal cortex (PPC) have been well-described (Johnston & Everling, 2011; Munoz & Everling, 2004; Paré & Dorris, 2011). In the anti-saccade task (where subjects must look away from a peripheral visual stimulus (Hallett, 1978)) DLPFC is thought to be critical to establishing the appropriate task set and preventing an automatic saccade to the stimulus; FEF is thought to be critical to voluntary saccade programming, and to “preparatory set”; and FEF, along with PPC are thought to be critical to the visuo-motor transformations to develop a saccade “vector” to the opposite direction of a stimulus (Connolly, Goodale, Menon, & Munoz, 2002; Leigh & Kennard, 2004; Munoz & Everling, 2004).

Evidence shows how DLPFC, FEF and PPC interact as part of a distributed system. For instance, TMS to *either* DLPFC, FEF (or supplementary eye fields) during saccade programming prolonged reaction times, suggesting that “preparatory set” was distributed between all three nodes (Nagel et al., 2008). Magnetoencephalography (MEG) and fMRI showed that FEF and PPC are both involved in the attentional aspects of the anti-saccade “vector” (Medendorp, Goltz, & Vilis, 2005; Moon et al., 2007), and TMS to FEF or PPC produces hypometric anti-saccades (Cameron, Riddle, & D’Esposito, 2015; Jaun-Frutiger, Cazzoli, Müri, Bassetti, & Nyffeler, 2013; Nyffeler, Hartmann, Hess, & Müri, 2008). However, it is not possible to distinguish a difference in timing (even with MEG) between when an anti-saccade program is developed in the PPC compared to FEF (Moon et al., 2007) implying a distributed process.

We build on this knowledge to study the effects on behavior after a “double lesion” to this network in the right hemisphere. Shortly after applying continuous theta-burst stimulation (Huang, Edwards, Rounis, Bhatia, & Rothwell, 2005) to either right FEF or DLPFC, we measure the consequences of a second time-resolved perturbation to the circuit, in the form of a single TMS pulse to right PPC. This approach arbitrates between five hypotheses regarding consequences of the double lesion. In hypothesis A – “*Additive*”, the double lesion could produce an additive effect by concurrently impairing spatially separate nodes that provide critical, but computationally distinct functions, resulting in behavioral perturbations that are greater than the effect of <u>either</u> perturbation alone (Figure 1A). Alternatively, hypothesis B – “*Distributed”* pertains to the case where computations are performed by a distributed system at the network-level, so a single lesion to either node should perturb behavior as much as the double lesion (Price, Hope, & Seghier, 2017) (Figure 1B). In hypothesis C – “*Compensatory*” - distal nodes could compensate for the perturbation, which would predict greater effects from the double lesion compared to the cTBS lesion alone (Figure 1C), because the second lesion impairs a region that has become more important functionally *because* of the first (cTBS) lesion. In hypothesis D – “*Spreading*” the effects from cTBS spread trans-synaptically to other portions of the network (Ko et al., 2008), predicting greater effects from the double lesion than to the single pulse lesion alone (Figure 1D). Finally, in “*Boosting”*, additional regions in the network could provide homeostatic compensation, which would manifest as a perplexing boost to performance following cTBS (alone), and which could reduce or prevent the impairment from additional TMS perturbations (Figure 1E). (The difference between this and the Compensatory hypothesis, is that there is the perplexing boost to performance after cTBS).

**Figure 1:**
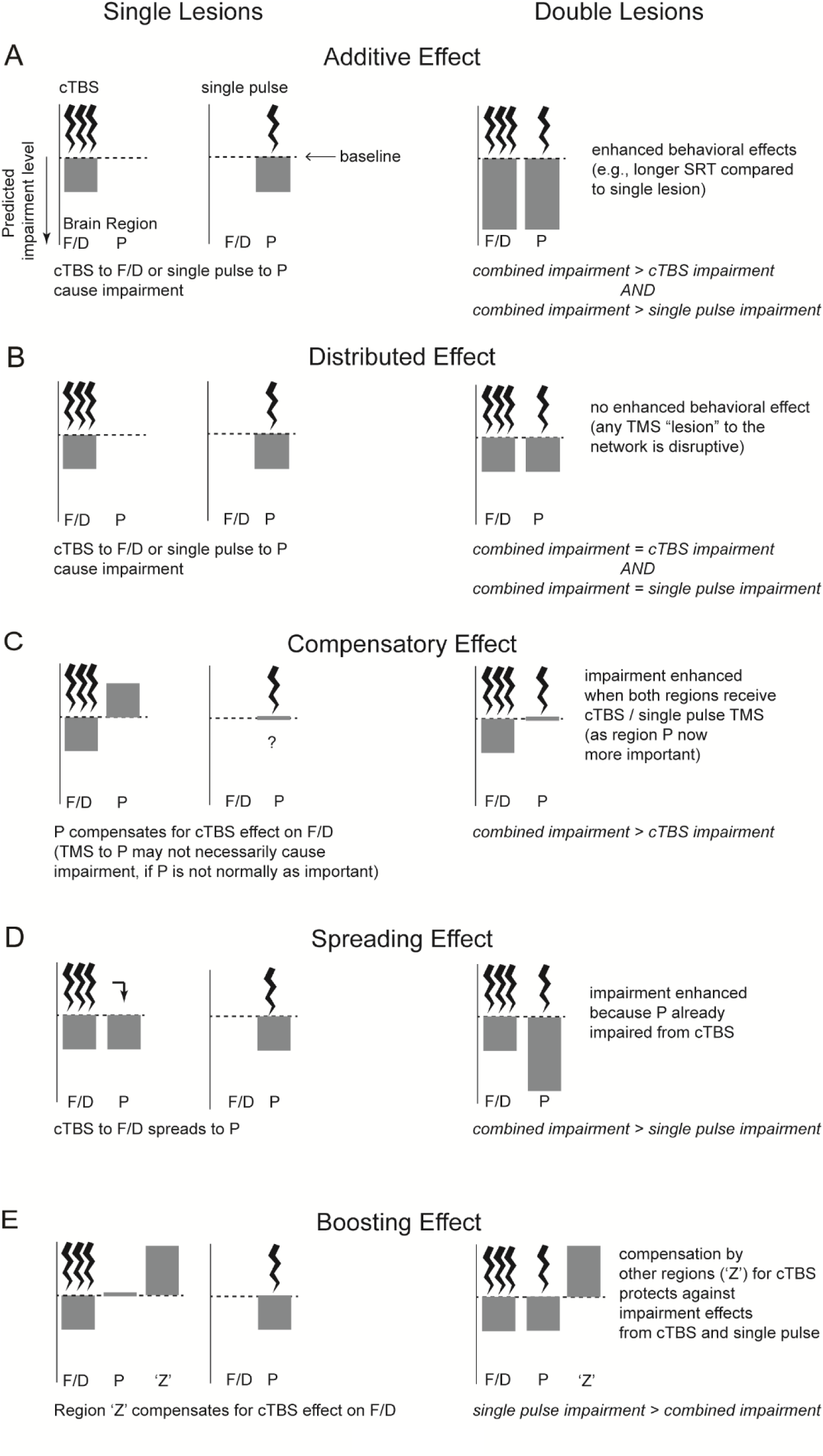
Hypotheses for the effects of TMS “lesions” to two oculomotor network nodes (e.g., F: frontal eye fields / D, dorsolateral prefrontal cortex; and P, posterior parietal cortex) in the same hemisphere. A) “Additive”: additive impairment from a double lesion compared to a single lesion to either node. B) “Distributed”: no additive effects (a single lesion to the network is disruptive). C) “Compensatory”: compensatory effect from second node that became more important. D) “Spreading”: enhanced effect due to cTBS spreading through the network to influence the second node. E) “Boosting”: additional network regions (region ‘Z’) provide sources of compensation after cTBS leading to a boost to performance.

To discriminate between those hypotheses, we first used functional magnetic resonance imagining (fMRI) to localize right DLPFC, FEF and PPC in individual human subjects performing an anti-saccade task. These regions were then used for targeting subject-specific TMS interventions while participants performed the same task outside the scanner. Performance (percentage correct direction), reaction times, and saccade amplitudes were assessed using Bayesian t-tests, useful to provide statistical evidence in favor or against enhanced effects from the double-compared to single-TMS lesions.

## Materials and methods

### Participants

The study was approved by the local ethics committee (Commissie Mensgebonden Onderzoek, Arnhem-Nijmegen) and written informed consent was obtained from the participants in accordance with the Declaration of Helsinki. A total of 27 healthy right-handed, young-adult, human subjects were recruited for 4 sessions approximately 1 week apart. 3 subjects were excluded for failure to provide useable eye-tracking data on all TMS sessions, and one subject had error rates on anti-saccade trials exceeding 90% (greater than 3 times the standard deviation), so was excluded resulting in a sample size of 24 participants (mean ± SE, age 23 ± 2 years, 11 male).

### Detailed procedure

#### Session 1

Participants were screened for contraindications related to fMRI, and to single-pulse TMS and cTBS according to common safety guidelines (Oberman, Edwards, Eldaief, & Pascual-Leone, 2011; Rossi, Hallett, Rossini, & Pascual-Leone, 2009). Resting and active motor thresholds were established for the first dorsal interosseous (FDI) muscle of the subject’s right-hand using electromyography (EMG). TMS was applied using a hand-held bi-phasic figure-eight coil with a 75 mm outer winding diameter (MagVenture, Denmark), connected to a MagProX100 system (MagVenture). Coil orientation was chosen to induce a posterior-anterior electrical field in the brain (45° from the mid-sagittal axis).

Subjects performed 5 runs of an interleaved pro-(look towards)/anti-(look away) saccade task to identify the cortical regions of interest (Figure 2B). An interleaved task was utilized as evidence suggests an important role for DLPFC (Everling & DeSouza, 2005; Johnston, Koval, Lomber, & Everling, 2014) as well as for FEF (DeSouza, Menon, & Everling, 2003) in task or “preparatory set” and thus could not simply default to an anti-saccade task set on each trial. Two target positions (13° or 9°) in the left and right direction were included so that subjects would have to rely on spatial information to calculate the saccade vector. In this way, we could be sure that the paradigm required DLPFC, FEF and PPC processes.

**Figure 2:**
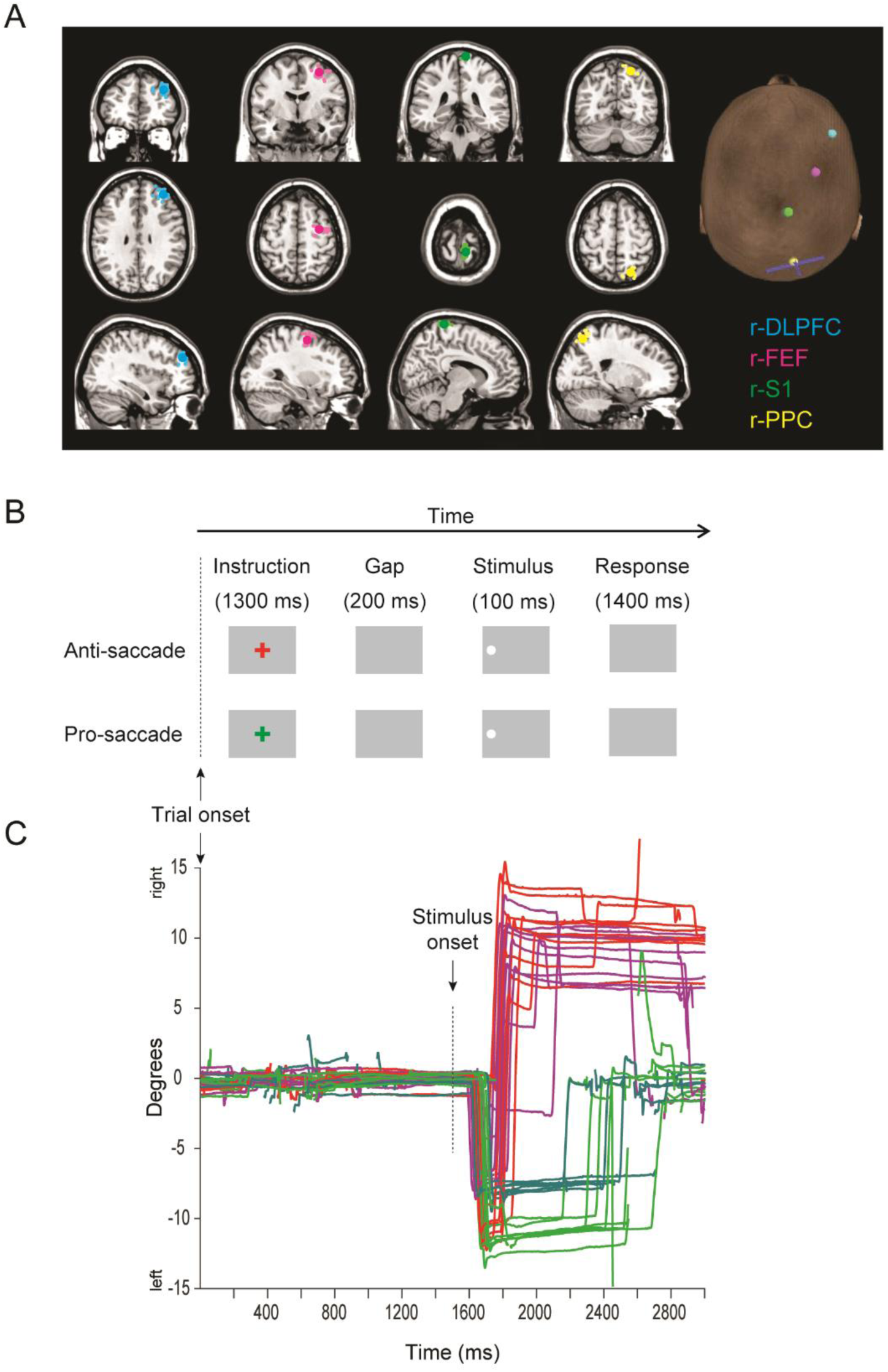
A) MRI images: illustration of coil placement over right dorsolateral prefrontal cortex (r-DLPFC), right frontal eye fields (r-FEF), right primary sensory cortex (r-S1), and right posterior parietal cortex (r-PPC) on an SPM single-subject anatomical template. Mean coordinates are shown as large bright dots, and individual subject coordinates are shown as faint dots. Right: scalp “entry” points for TMS stimulation for a representative subject, showing also a representation of the coil orientation over right PPC (handle of coil = base of ‘T’ shape). B) Paradigm and stimulus timings shown for representative anti-saccade and pro-saccade trials, where the target stimulus was on the left side. C) Illustrations of raw eye-traces from a representative subject in one run (subject 22841) with respect to stimuli on the left side. For 13° stimuli, red illustrates anti-saccades and green illustrates pro-saccades; for 9° stimuli, magenta illustrates anti-saccades, and turquoise illustrates pro-saccades. This subject made a high proportion of direction errors on anti-saccade trials in this run, indicated by the reversals of direction. Blinks are shown as gaps in the traces.

#### Detailed fMRI procedure

Functional MRI scans were obtained with a 3 Tesla MRI scanner (Skyra, Siemens Medical Systems Erlangen, Germany) using a 32-channel head coil. The functional images were acquired with multiband sequence (acceleration factor = 3, repetition time (TR) = 1000 ms, echo time (TE) = 30 ms, flip angle= 60°). Each volume consisted of 33 slices, with a distance of 17% and a thickness of 3 mm. The voxel resolution was 3.5 × 3.5 × 3.0 mm, FoV in the read direction of 224 mm and FoV in the phase direction of 100%. Two volumes were discarded from each functional run, to account for scanner steady state equilibrium, leading to a total of 339 volumes per run. The anatomical images were acquired with a MPRAGE sequence (repetition time (TR) = 2300 ms, echo time (TE) = 3.9 ms, voxel size = 1 × 1 × 1 mm). In total, 192 images were obtained for each participant. During the scan, participants lay in a supine position and their head was stabilized using soft cushions.

Imaging data were analyzed with SPM8 (Wellcome Trust Centre for Cognitive Neuroimaging, London, UK). At the single-subject level, the data were realigned to the first volume of each run using six rigid body transformations (3 translations and 3 rotations). The images were then coregistered to the individual structural T1 and spatial smoothing was performed by means of an 8-mm full-width half-maximum (FWHM) Gaussian kernel. A first-level analysis was performed by specifying a general linear model with regressors for each condition (fixation trials were not modeled however). Motion parameters (3 translations, 3 rotations) were included as nuisance regressors.

A contrast of anti-saccade trials against baseline was computed to define 5 mm ROIs centered on locations of peak activation on each subject anatomical scan, using a t-contrast at P < 0.001 (uncorrected). Table 1 provides the Montreal Neurological Institute (MNI) coordinates of these ROIs, and their distances to the scalp as derived from Localite TMS Navigation software 2.2 (Localite, Germany). Figure 2A illustrates the coordinates on a canonical T1 scan. Right DLPFC (r-DLPFC) was defined as peak fMRI anti-saccade activity surrounding the middle frontal gyrus, anterior to the ventricles. Right FEF was defined as peak activity in the precentral sulcus (selecting medial peaks if lateral peaks were also present, to relate more to anti-saccade processes (Neggers et al., 2012)). Right PPC was defined as peak activity in the intraparietal sulcus, selecting peaks in more medial clusters if more than one was present. Finally, right S1 (the control region) was localized anatomically for each participant, as the most superior extent of the postcentral gyrus, located on average 9 ± 2 mm lateral to the longitudinal fissure to avoid lateral proprioceptive eye-position signals (Balslev, Albert, & Miall, 2011; Zhang, Wang, & Goldberg, 2008) (Table 1, Figure 2A).

**Table 1:**
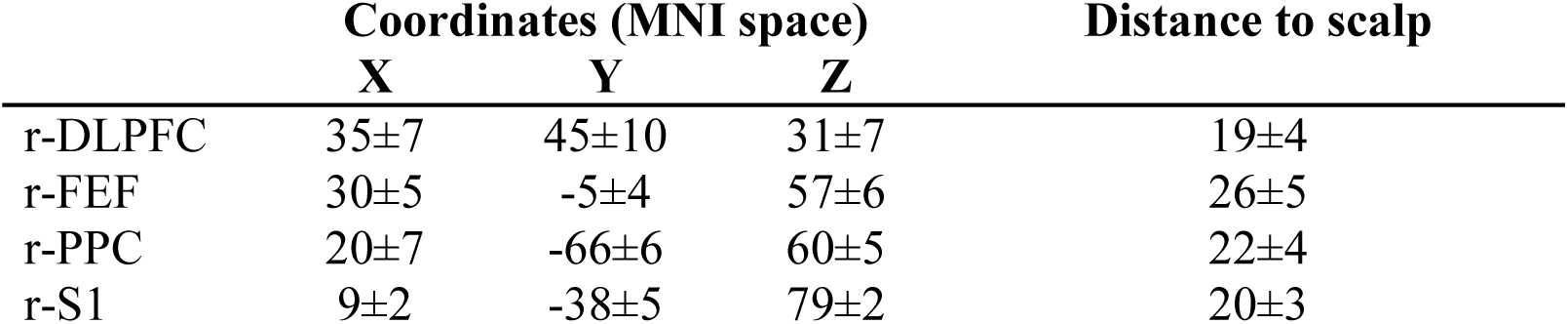
Regions Of Interest (ROI) information (average ± standard deviations (mm))

|  | Coordinates (MNI space) |  |  | Distance to scalp |
| --- | --- | --- | --- | --- |
|  | X | Y | Z |  |
| r-DLPFC | 35±7 | 45±10 | 31±7 | 19±4 |
| r-FEF | 30±5 | -5±4 | 57±6 | 26±5 |
| r-PPC | 20±7 | -66±6 | 60±5 | 22±4 |
| r-S1 | 9±2 | -38±5 | 79±2 | 20±3 |

#### Session 2-4

cTBS was applied to r-DLPFC, r-FEF or r-S1 prior to performing the task on three separate sessions, counterbalanced for order. cTBS was applied to FEF or to DLPFC because we wished to assess double lesion effects across two nodes which are both linked to PPC, but where one (FEF) is thought to have a more direct link in visuo-motor processes (Leigh & Kennard, 2004; Munoz & Everling, 2004) and in network interactions described in the resting state (Corbetta, Kincade, Lewis, Snyder, & Sapir, 2005; He et al., 2007; Vossel, Geng, & Fink, 2014). cTBS was delivered with a posterior-anterior direction of the electric field induced in the brain, with the handle pointed backwards at approximately 30° to the sagittal plane. In this way the outer windings of the TMS coil did not overlap the other ROIs. TMS coil alignment was achieved using Localite and a subject-specific anatomical scan.

The parameters for cTBS were identical to those described by Huang and colleagues (2005) consisting of 50 Hz triplets repeated at 5 Hz over a period of 40 s (Huang et al., 2005). Stimulation intensity for cTBS was defined as 80% of the active motor threshold (AMT), defined as peak-to-peak MEP amplitudes exceeding 200 μV on 5 out of 10 trials, while subjects maintained voluntary contraction of approximately 10%. Stimulation intensity for single pulse TMS to PPC was set at 110% of the resting motor threshold (RMT), defined as peak-to-peak MEP amplitudes of 50 μV on 5 of 10 trials. 40 s of cTBS (at 80% of active motor threshold) has effects lasting approximately 50 minutes (Wischnewski & Schutter, 2015), providing sufficient time to test the influence of the PPC pulse.

#### Eye Tracking and Task

The position of the right eye was recorded using an infrared Eyelink 1000 eye tracker (SR Research, Ottawa, Canada) with a 1000 Hz sampling rate. A 9-point calibration was carried out and a drift correction point was used as the inter-trial fixation point. Saccades were identified by a horizontal deflection (3 X standard deviations of the baseline velocity) and duration between 15 and 150 ms. The camera was positioned under the stimulus screen, approximately 60 cm away from the eyes of the participant, who sat precisely at 70 cm from a wide-angle LCD screen (with central presentation zone set at 4:3, 1024 × 768 resolution).

Subjects performed the same task (Figure 2B) as in the fMRI. Representative eye-traces from a single subject is shown in Figure 2C. In each run, there were 72 trials, of which 48 contained a TMS pulse presented to PPC at a random interval between 30 and 300 ms after onset of the peripheral stimulus (described in Data Analysis). The first run commenced 10 minutes after cTBS, and was analyzed up to 50 minutes after cTBS to capture the same cTBS effects on each session. Subjects were asked to perform 5 runs, each taking approximately 8 minutes including drift corrections and breaks, meaning that for each condition of interest (task, direction) there were 30 trials without the single pulse (“*pulse absent*”), and 60 trials containing the single pulse.

### Data analysis

Data was analyzed with custom MATLAB v11 programs (The MathWorks Inc., Natick, MA). Valid trials consisting of correct and incorrect directions were separated from invalid trials, consisting of saccade reaction times (SRTs) < 90 ms (anticipatory errors), slower than 1000 ms, and trials where the TMS pulse to PPC occurred after saccade onset. Three behavioral parameters of interest were analyzed: amplitude of the primary saccade, percentage correct direction, and saccade reaction time (SRT).

We first set a division between an “Early” and “Late” pulse time bin as follows: using the pulse absent trials, we collected the SRTs across subjects for correctly performed anti-saccades, and for direction errors on anti-saccades for each cTBS session separately, and plotted these data in 10 ms bin histograms (Figure 3). A binomial test revealed the first bin (black arrows, Figure 3) where the two trial types were no longer significantly different than chance (50 %); these bins occurred at 150 ms for the S1 cTBS and DLPFC cTBS sessions, and at 160 ms for the FEF-cTBS session. This method approximates the division between visually triggered “express” pro-saccades, and voluntary saccades (Munoz & Everling, 2004), and is important to approximate when the PPC pulse would have greater influences during visual processing rather than motor programming components of an anti-saccade, which are in different directions.

**Figure 3:**
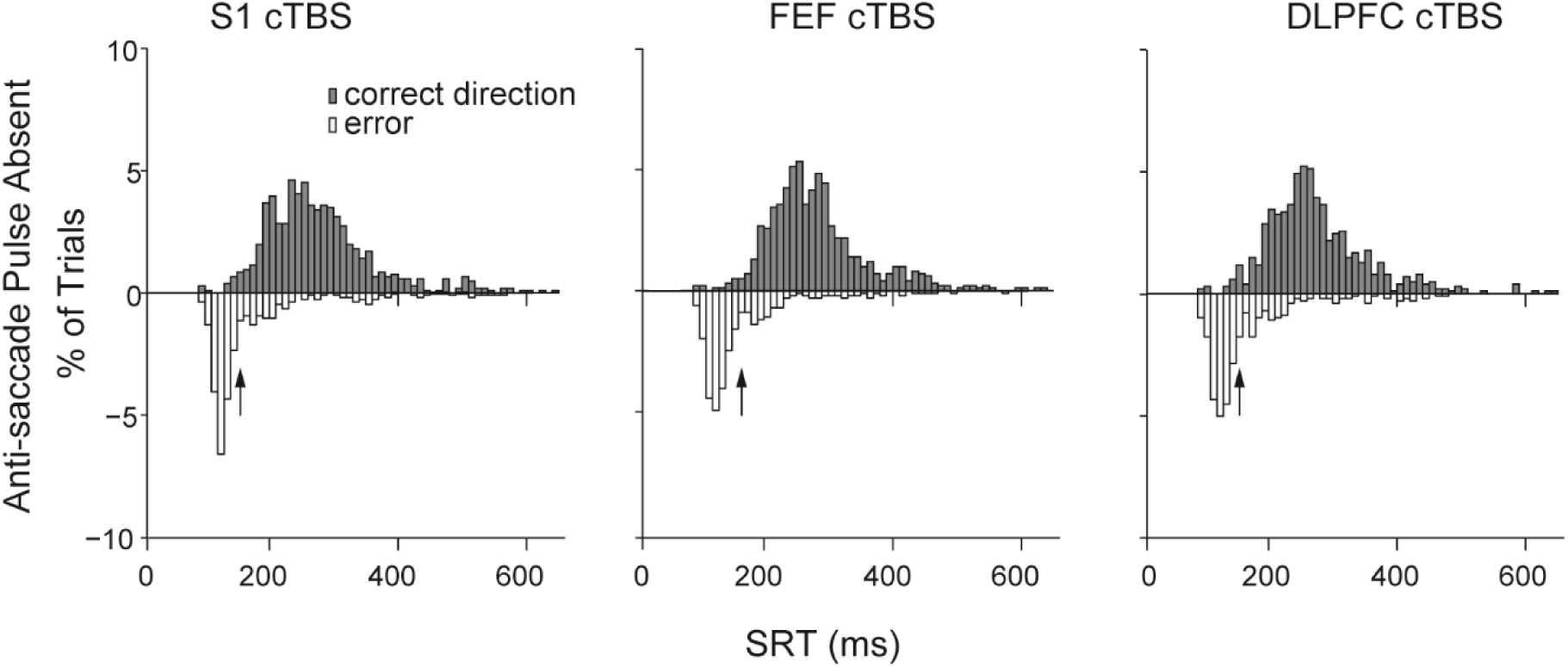
Derivation of the Early and Late PPC pulse bins based on anti-saccade reaction times. Reaction time distributions were calculated for correct and direction error anti-saccades in PPC Pulse Absent trials on each cTBS session. A binomial sign test was performed compared the distributions, and arrows indicate the first reaction time bin where the two distributions were no longer significantly different. This value was taken as the boundary for Early and Late PPC pulses.

Next, we performed a repeated measures ANOVA using pulse absent trials to determine if there were significant interactions between the site of cTBS and stimulus eccentricity for amplitudes. However, no interactions with cTBS Site and Eccentricity were significant, *F*(2,44) < 1.75, *P* > 0.19, so we collapsed across eccentricity.

To directly assess our five network hypotheses regarding the combined effects from cTBS and the PPC pulse (Figure 1), we performed Bayesian paired-sample t-tests in JASP (JASP Team, 2017) (Figures 4-7, brackets). A Bayes Factor (BF_10_) indicates the evidence for the alternative hypothesis relative to the null hypothesis given the data. Our tests were focused first on situations where the “double lesion” produces impairments that were greater than the single lesions; thus, the Bayes Factor (BF_10_) here indicates whether the *combined effects* were greater than the individual effects from cTBS alone, or from PPC TMS alone). For amplitude and percent correct, lower values are indicative of greater impairments: therefore, the alternative hypothesis for BF_10_ is that the difference of the combined effect minus the single lesion effect was less than 0, and the null hypothesis would be that this difference is not less than zero. For reaction times, higher values are indicative of an impairment (slower latency), so the alternative hypothesis is that the combined effect minus the single lesion effect is greater than zero (and the null hypothesis is that it is not greater than zero). Note, however, that strong evidence from these tests for the null hypothesis (not less than zero) could be driven by a difference in the opposite direction. When such “strong” evidence was found (BF_10_ < 0.1) (Jeffreys, 1961; Wetzels et al., 2011), we subsequently performed tests in the opposite direction to determine if the effect of the single lesion was greater than that of the double lesion.

**Figure 4:**
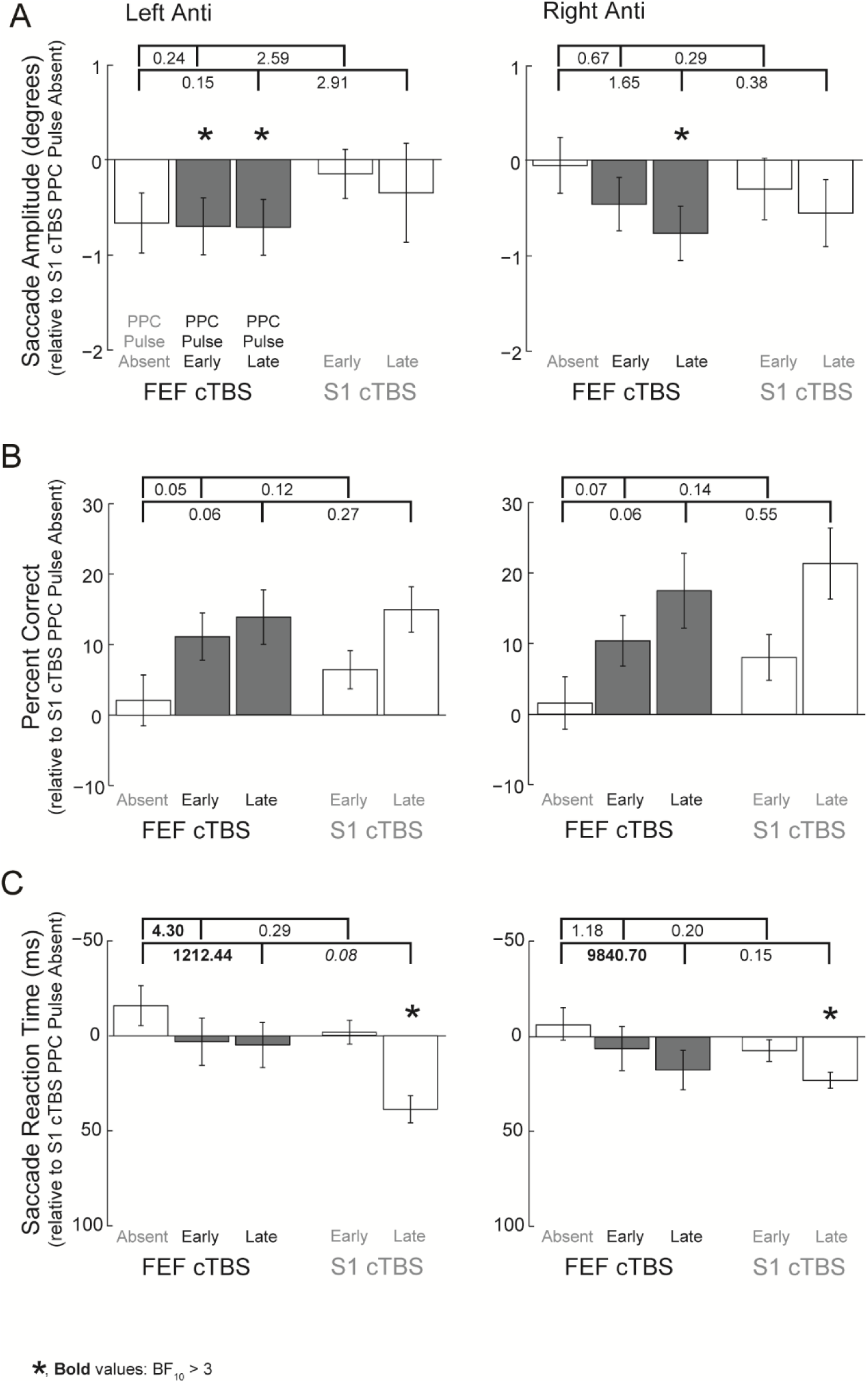
Effects on left and right anti-saccades when the double lesion involved FEF cTBS and PPC TMS is compared to the single lesion conditions. All data is normalized to the cTBS control condition (cTBS to S1, no PPC pulses). Error bars represent standard error of the mean across subjects (N=23), and dark grey represents the double lesion conditions. Values between brackets indicate the Bayes Factor evidence for the alternative hypothesis that the combined effects from the double lesion resulted in a greater impairment (more negative values, note the Y axis is reversed for reaction times) compared to the effects of the single lesions. Values > 3 provide substantial evidence for the alternative hypothesis that the combined effects resulted in a greater impairment than the single lesion effects. Asterisks show the results from Bayesian one-sample t-tests for evidence that the values are < 0 where BF_10_ > 3.

We report evidence for behavioral impairments that meet or exceed “*substantial*” (BF_10_ > 3) (Jeffreys, 1961; Wetzels et al., 2011). Between 0.33 and 3, the evidence is considered only “*anecdotal*”, and in relation to P-values, it was shown that approximately 70% of “positive” results from 855 tests falling in the interval between *P* < 0.01 to *P* < 0.05 corresponded to only “anecdotal” evidence (Wetzels et al., 2011). Therefore, our boundary criteria of “substantial” is conservative in relation to typical P-values.

Tests for each individual trial type compared to the control condition (S1 cTBS, PPC Pulse Absent) were also conducted using Bayesian one-sample t-tests in JASP to confirm if the individual lesions themselves caused impairments. Here, the BF_10_ indicates the relative likelihood that cTBS or single pulse TMS impaired behavior compared to the null hypothesis that the effects were equal to the control condition. The values for these tests are listed in Tables 2-5, and are illustrated as asterisks in Figure 4-7 when substantial.

**Table 2.** Bayes Factors for the alternative (impairment) vs. null (no impairment) hypothesis (BF_10_) for left and right anti-saccade trials relative to control cTBS

| **Left**<br>**Anti** | cTBS<br>site | PPC<br>pulse | $BF_{10}$ | **Right**<br>**Anti** | cTBS<br>site | PPC<br>pulse | $BF_{10}$ |
| --- | --- | --- | --- | --- | --- | --- | --- |
| **A) Amplitude** |  |  |  |  |  |  |  |
|  | FEF | Absent | 2.69 |  | FEF | Absent | 0.26 |
|  |  | Early | **4.09** |  |  | Early | 1.31 |
|  |  | Late | **4.58** |  |  | Late | **7.60** |
|  | S1 | Early | 0.35 |  | S1 | Early | 0.52 |
|  |  | Late | 0.39 |  |  | Late | 1.21 |
| **B) Percent Correct** |  |  |  |  |  |  |  |
|  | FEF | Absent | 0.15 |  | FEF | Absent | 0.16 |
|  |  | Early | 0.06 |  |  | Early | 0.07 |
|  |  | Late | 0.06 |  |  | Late | 0.06 |
|  | S1 | Early | 0.08 |  | S1 | Early | 0.07 |
|  |  | Late | 0.05 |  |  | Late | 0.06 |
| **C) Saccade Reaction Time** |  |  |  |  |  |  |  |
|  | FEF | Absent | 0.59 |  | FEF | Absent | 0.29 |
|  |  | Early | 0.23 |  |  | Early | 0.25 |
|  |  | Late | 0.24 |  |  | Late | 0.74 |
|  | S1 | Early | 0.22 |  | S1 | Early | 0.45 |
|  |  | Late | **1149.80** |  |  | Late | **1309.96** |
| **Bold values: $BF_{10} > 3$** |  |  |  |  |  |  |  |

## Results and Discussion

### FEF cTBS vs control cTBS conditions: anti-saccades

#### Saccade amplitude

There was substantial evidence that FEF cTBS caused impairments in leftward anti-saccade amplitudes for conditions also involving PPC pulses, and for rightwards anti-saccades for conditions involving the late PPC pulse (Table 2A, BF_10_ > 3). There was not substantial evidence that the PPC pulse *on its own* produced an impairment, and there was also *not* substantial evidence (Figure 4A, brackets, all BF_10_ ≤ 2.91) to indicate enhanced impairments from the double lesion condition compared to either single lesion condition.

#### Percentage correct direction

There was not substantial evidence that anti-saccades were impaired by either form of TMS; in fact, *strong* evidence towards the null hypothesis was found for conditions with the PPC pulse (Table 2B, BF_10_ < 0.1). (Bayesian t-tests performed in the opposite direction revealed substantial or greater evidence (BF_10_ > 3) for a performance *benefit* from the PPC pulses). Similarly, there was strong evidence that there were *not* enhanced impairments from the double lesion compared to either single lesion (Figure 4B).

#### Saccade Reaction Times (SRT)

For SRT, “*decisive”* (Wetzels et al., 2011) evidence for impairments were observed for conditions with the late PPC pulse alone, but not for those following FEF cTBS (Table 2C). Strong evidence was found that FEF cTBS plus a late PPC pulse did result in enhanced impairments relative to FEF cTBS alone (and substantial evidence was found for an enhanced impairment for the early PPC pulse for leftwards anti-saccades) (Figure 4C). However, strong evidence was found that impairments for leftwards anti-saccades were *not* greater when the late PPC pulse followed FEF cTBS compared to when it was alone (Figure 4C, BF = 0.08, italicized): when tested in the reverse direction, there was substantial evidence that the impairment after the late PPC pulse *alone* was greater than after FEF cTBS, BF_10_ = 4.12.

#### Interpretation

We did not find evidence to suggest Hypothesis A (an “additive” effect) from the double lesion across any of the saccade behaviors. Substantial evidence showed impairments to anti-saccade amplitude following FEF cTBS when PPC pulses were present; however, because there was not substantial evidence that PPC pulses on their own caused impairments, nor were the effects enhanced relative to those following FEF cTBS, we conclude that cTBS to FEF was consequential to anti-saccade amplitudes suggesting a “distributed” effect (Hypothesis B).

For saccade reaction times, we found evidence for enhanced impairments from the double lesion compared to FEF cTBS on its own. We do not believe this indicates compensation by PPC, because these conditions (FEF cTBS plus PPC pulse) did not reveal substantial evidence for impairment on their own. In fact, the late PPC pulses on their own produced impairments that were *greater* than the double lesion for contralateral anti-saccades (Hypothesis E). We conclude therefore that later PPC pulses were disruptive to the motor component of the anti-saccade. Following FEF cTBS however, a compensatory mechanism might be revealed by other network structures (e.g., the superior colliculus, SC) which aid in anti-saccade generation. One possibility is that after FEF cTBS, there is compensation by DLPFC-colliculus projections to contralateral SC saccade neurons (Everling & Johnston, 2013), reducing the disruptive effect from a PPC pulse on the same network structures. This is sensible, considering the PPC pulses also produced substantial anti-saccade *performance* benefits, and human EEG evidence has shown that the posterior parietal/occipital cortex is involved in triggering “express” pro-saccades (Hamm, Dyckman, Ethridge, McDowell, & Clementz, 2010), possibly by a cortical-SC mechanism (Chen, Liu, Wei, & Zhang, 2013; Watanabe, Hirai, Marino, & Cameron, 2010). A PPC pulse could thus disrupt the bias towards stimulus-driven saccades thus indirectly facilitating anti-saccade performance.

However, these effects could also be driven by the auditory/or somatosensory influence of the pulse (Duecker, de Graaf, Jacobs, & Sack, 2013; Duecker & Sack, 2013), which could engage a startle-like reflex that inhibits ongoing motor commands, by acting also on the SC or brain stem saccade generator circuits (Xu-Wilson, Tian, Shadmehr, & Zee, 2011) (perhaps with less of a consequence in cases of compensation). As the goal of this study was to compare hypotheses regarding the double vs single lesions situations, the interesting comparisons are those between the PPC pulses following control versus verum cTBS, which both have the same auditory/somatosensory influences of the PPC pulse.

### DLPFC cTBS vs control cTBS conditions: anti-saccades

#### Saccade amplitude

There was substantial evidence for impairments to anti-saccades after DLPFC cTBS in conditions involving the late PPC pulse, and for DLPFC cTBS alone for leftward anti-saccades (Table 3A). Strong evidence was found for an *enhanced* impairment from the combined lesion effects for rightward anti-saccades after the late pulse relative to the DLPFC cTBS alone (BF_10_ = 325.22), but this was not found compared to the effects of the late PPC pulse alone (BF_10_ = 0.75) (Figure 5A).

**Table 3:** Bayes Factors for the alternative (impairment) vs. null (no impairment) hypothesis (BF_10_) for left and right anti-saccade trials relative to control cTBS. (The effect of the PPC pulse relative to control cTBS is shown in duplication as in Table 3)

| Left<br>Anti | cTBS<br>site | PPC<br>pulse | $BF_{10}$ | Right<br>Anti | cTBS<br>site | PPC<br>pulse | $BF_{10}$ |
| --- | --- | --- | --- | --- | --- | --- | --- |
| **A) Amplitude** |  |  |  |  |  |  |  |
|  | DLPFC | Absent | **4.21** |  | DLPFC | Absent | 0.33 |
|  |  | Early | 0.55 |  |  | Early | 0.98 |
|  |  | Late | **4.84** |  |  | Late | **8.84** |
|  | S1 | Early | 0.35 |  | S1 | Early | 0.52 |
|  |  | Late | 0.39 |  |  | Late | 1.21 |
| **B) Percent Correct** |  |  |  |  |  |  |  |
|  | DLPFC | Absent | 0.27 |  | DLPFC | Absent | 0.16 |
|  |  | Early | 0.14 |  |  | Early | 0.09 |
|  |  | Late | 0.05 |  |  | Late | 0.05 |
|  | S1 | Early | 0.08 |  | S1 | Early | 0.07 |
|  |  | Late | 0.05 |  |  | Late | 0.06 |
| **C) Saccade Reaction Time** |  |  |  |  |  |  |  |
|  | DLPFC | Absent | 0.24 |  | DLPFC | Absent | 0.40 |
|  |  | Early | 0.46 |  |  | Early | 0.24 |
|  |  | Late | 1.30 |  |  | Late | **16.96** |
|  | S1 | Early | 0.22 |  | S1 | Early | 0.45 |
|  |  | Late | **1149.80** |  |  | Late | **1309.96** |
| **Bold values: $BF_{10} > 3$** |  |  |  |  |  |  |  |

**Figure 5:**
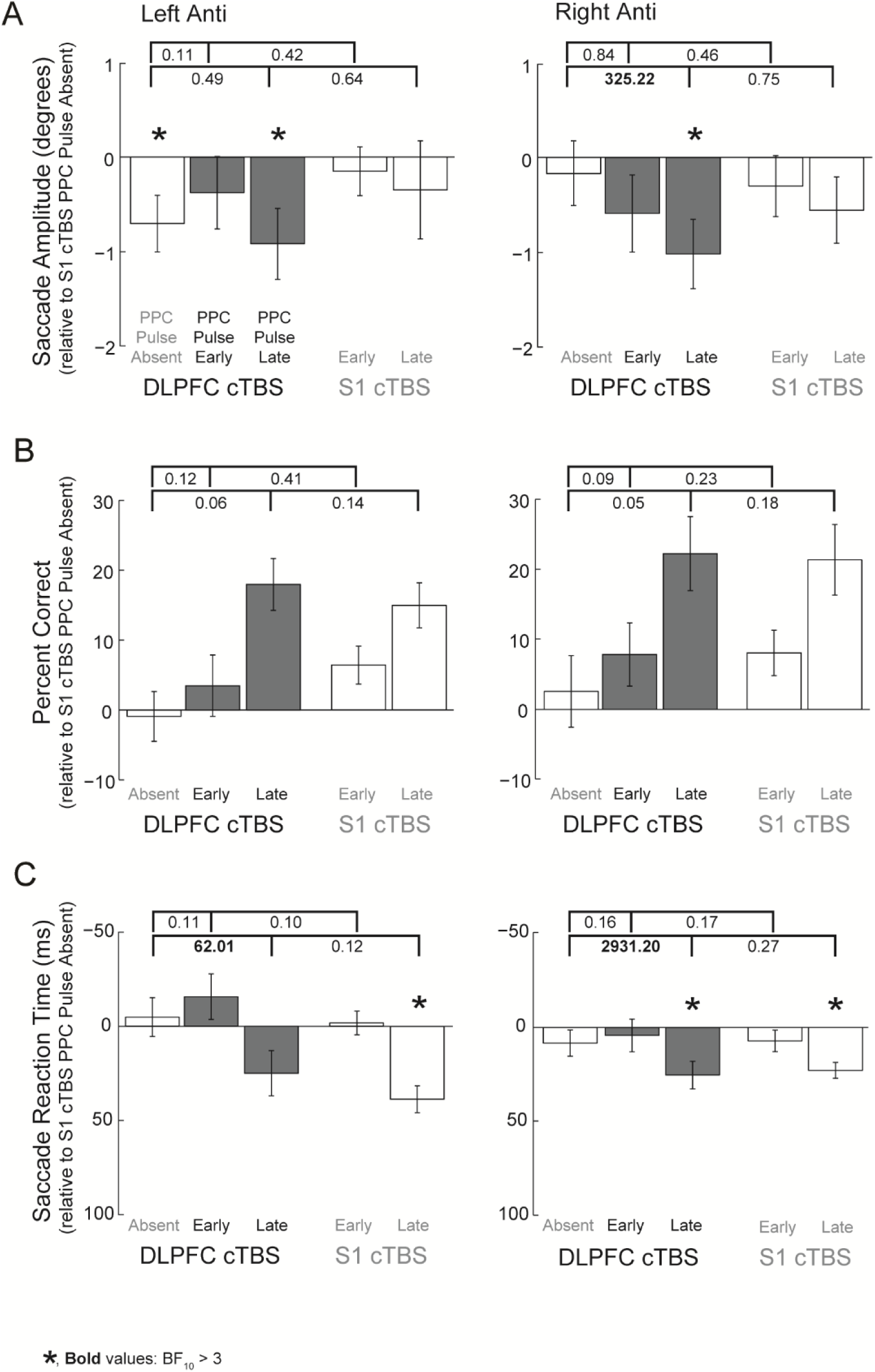
Effects on left and right anti-saccades when the double lesion involving DLPFC cTBS and PPC TMS is compared to the single lesion conditions. PPC pulse conditions relative to S1 cTBS are shown in duplication as in Figure 4, and conventions are as in Figure 4.

#### Percentage correct direction

There was no evidence that anti-saccades were impaired by DLPFC cTBS, with, or without, the PPC pulse (Table 3B). (Bayesian t-tests revealed strong evidence for anti-saccade *benefits* to performance following DLPFC cTBS and late PPC pulses). There was also no evidence for enhanced impairment from a double compared to single lesion (Figure 5B).

#### Saccade Reaction Times (SRT)

There was strong evidence for impaired reaction times at the late pulse time following DLPFC cTBS for right anti-saccades (Table 3C), and there was strong evidence that the combined effects of DLPFC cTBS and a late PPC pulse resulted in enhanced impairments relative to DLPFC cTBS alone (Figure 5C), but there was no evidence for greater impairment in comparison to the PPC pulse.

#### Interpretation

We found strong evidence for *compensation* by PPC (Hypothesis C) following DLPFC cTBS for ipsilateral (rightward) anti-saccade amplitudes, but not substantial evidence for an additive effect (Hypothesis A). Importantly, there was not substantial evidence that the PPC pulses alone produced an impairment. This is consistent with a compensatory hypothesis, in that the second lesion impairs a node which has as a greater contribution (Hartwigsen et al., 2016; Sack et al., 2005). We note that these effects were lateralized, as compensation was only seen in this ipsilateral direction. cTBS to DLPFC alone produced impairments in the contralateral direction, suggesting that DLPFC lesions were more consequential for contralateral anti-saccades.

In a previous study, we did not find significant impairments in saccade amplitudes following (left) DLPFC cTBS (Cameron et al., 2015), and it has also been concluded by one lesion study that DLPFC was not necessary for performing the spatial calculations in a memory-guided saccade task (Mackey, Devinsky, Doyle, Meager, & Curtis, 2016). However, another study did show DLPFC lesions resulted in higher variability in memory-guided saccade endpoints, with non-significant reductions in amplitudes (Pierrot-Deseilligny et al., 2003). Such mixed findings lend support to a hypothesis that the spatial calculations for anti-saccades are performed by a distributed process. We can only speculate that compensation occurs in some circumstances depending on the particular task.

As with FEF cTBS, there was no evidence that any of the conditions impaired percentage correct direction, but there was evidence for enhanced SRT impairments from the combined double lesion compared to DLPFC cTBS alone. As addressed previously, the late PPC pulse impaired SRT on its own, suggesting the effects are more related to that of the PPC pulse.

### FEF cTBS vs control cTBS conditions: pro-saccades

#### Saccade amplitude

Table 4A and Figure 6A show that there was not substantial evidence for effects of either TMS condition on pro-saccade amplitudes.

**Table 4:** Bayes Factors for the alternative (impairment) vs. null (no impairment) hypothesis (BF_10_) for left and right pro-saccade trials relative to control cTBS

| **Left**<br>**Pro** | cTBS<br>site | PPC<br>pulse | $BF_{10}$ | **Right**<br>**Pro** | cTBS<br>site | PPC<br>pulse | $BF_{10}$ |
| --- | --- | --- | --- | --- | --- | --- | --- |
| **A) Amplitude** |  |  |  |  |  |  |  |
|  | FEF | Absent | 0.61 |  | FEF | Absent | 0.54 |
|  |  | Early | 0.31 |  |  | Early | 0.56 |
|  |  | Late | 0.87 |  |  | Late | 0.30 |
|  | S1 | Early | 0.11 |  | S1 | Early | 1.07 |
|  |  | Late | 0.30 |  |  | Late | 2.32 |
| **B) Percent Correct** |  |  |  |  |  |  |  |
|  | FEF | Absent | 0.12 |  | FEF | Absent | 1.16 |
|  |  | Early | 0.41 |  |  | Early | 0.22 |
|  |  | Late | 2.63 |  |  | Late | **4.53** |
|  | S1 | Early | 0.48 |  | S1 | Early | 0.24 |
|  |  | Late | 1.44 |  |  | Late | 1.19 |
| **C) Saccade Reaction Time** |  |  |  |  |  |  |  |
|  | FEF | Absent | 0.23 |  | FEF | Absent | 0.24 |
|  |  | Early | **7.47** |  |  | Early | 2.08 |
|  |  | Late | **1657.49** |  |  | Late | **730.84** |
|  | S1 | Early | **55.31** |  | S1 | Early | **25.74** |
|  |  | Late | **26318.61** |  |  | Late | **1082000** |
| **Bold values: $BF_{10} > 3$** |  |  |  |  |  |  |  |

**Figure 6:**
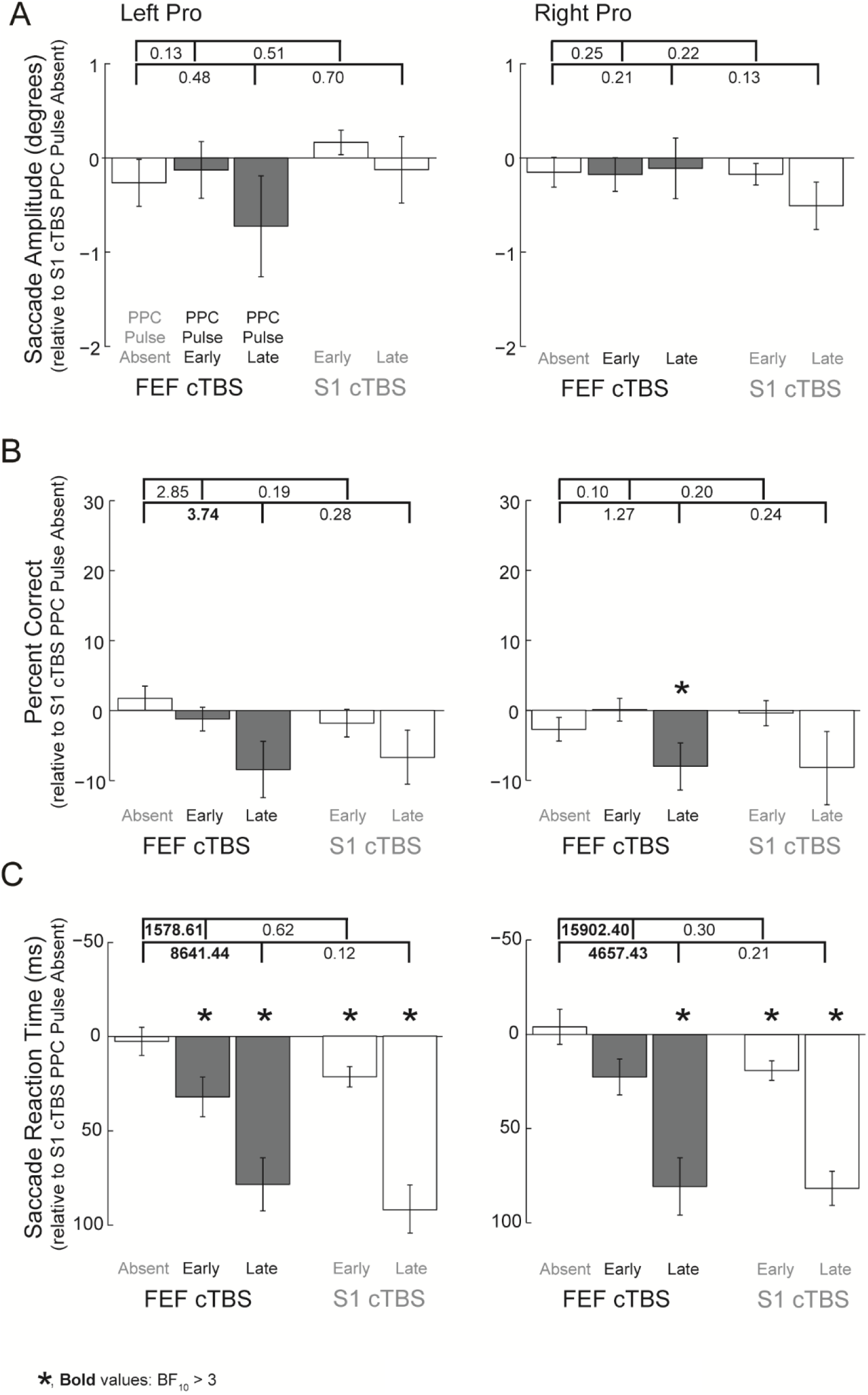
Effects on left and right pro-saccades when the double lesion involving FEF cTBS and PPC TMS is compared to the single lesion conditions. Conventions as in Figure 4.

#### Percentage correct direction

Substantial impairments were found for rightwards pro-saccades following FEF cTBS during trials with the addition of a late PPC pulse (Table 4B; BF_10_ = 4.53). There was also substantial evidence that the impairments to leftwards pro-saccades were greater following FEF cTBS when there was a late PPC pulse (Figure 6B; BF_10_ = 3.74) compared to FEF cTBS alone. There was not substantial evidence for other impairments.

#### Saccade Reaction Times (SRT)

Substantial or greater evidence for pro-saccade reaction time impairments was observed for 7/8 PPC pulse conditions (Table 4C). There was also strong evidence that the combined effects of FEF cTBS and PPC pulses resulted in enhanced impairments relative to FEF cTBS alone (Figure 6C), however, there was not evidence for an enhanced impairment over the PPC pulse effects alone.

#### Interpretation

There was no evidence to suggest that TMS to FEF or PPC impaired pro-saccade amplitudes, suggesting that other regions in a wider network are sufficient for the spatial calculations for a pro-saccade (Munoz & Schall, 2004). There were findings to suggest that the late PPC pulse following FEF cTBS impaired pro-saccade correct directions and substantially increased reaction times, suggesting a detrimental effect of the PPC pulse, possibly by impairing PPC-SC signals (as described previously). We acknowledge, however, that because we rejected trials when reaction time was less than the PPC pulse time, the outcome measures of the late PPC pulse are biased as coming from trials with a slower latency.

### DLPFC cTBS vs control cTBS conditions: pro-saccades

#### Saccade amplitude

There was not substantial evidence for any effects to pro-saccade amplitudes (Table 5A, Figure 7A).

**Table 5:** Bayes Factors for the alternative (impairment) vs. null (no impairment) hypothesis (BF_10_) for left and right pro-saccade trials relative to control cTBS (The effect of the PPC pulse relative to control cTBS is shown in duplication as in Table 4)

| Left<br>Pro | cTBS<br>site | PPC<br>pulse | $BF_{10}$ | Right<br>Pro | cTBS<br>site | PPC<br>pulse | $BF_{10}$ |
| --- | --- | --- | --- | --- | --- | --- | --- |
| **A) Amplitude** |  |  |  |  |  |  |  |
|  | DLPFC | Absent | 0.19 |  | DLPFC | Absent | 0.62 |
|  |  | Early | 0.24 |  |  | Early | 0.22 |
|  |  | Late | 0.15 |  |  | Late | 1.03 |
|  | S1 | Early | 0.11 |  | S1 | Early | 1.07 |
|  |  | Late | 0.30 |  |  | Late | 2.32 |
| **B) Percent Correct** |  |  |  |  |  |  |  |
|  | DLPFC | Absent | 0.22 |  | DLPFC | Absent | 0.17 |
|  |  | Early | 0.32 |  |  | Early | 0.21 |
|  |  | Late | 0.42 |  |  | Late | 2.84 |
|  | S1 | Early | 0.48 |  | S1 | Early | 0.24 |
|  |  | Late | 1.44 |  |  | Late | 1.19 |
| **C) Saccade Reaction Time** |  |  |  |  |  |  |  |
|  | DLPFC | Absent | 0.23 |  | DLPFC | Absent | 0.34 |
|  |  | Early | 1.86 |  |  | Early | 1.29 |
|  |  | Late | **2544.68** |  |  | Late | **1172000** |
|  | S1 | Early | **55.31** |  | S1 | Early | **25.74** |
|  |  | Late | **26318.61** |  |  | Late | **1082000** |
| **Bold values: $BF_{10} > 3$** |  |  |  |  |  |  |  |

**Figure 7:**
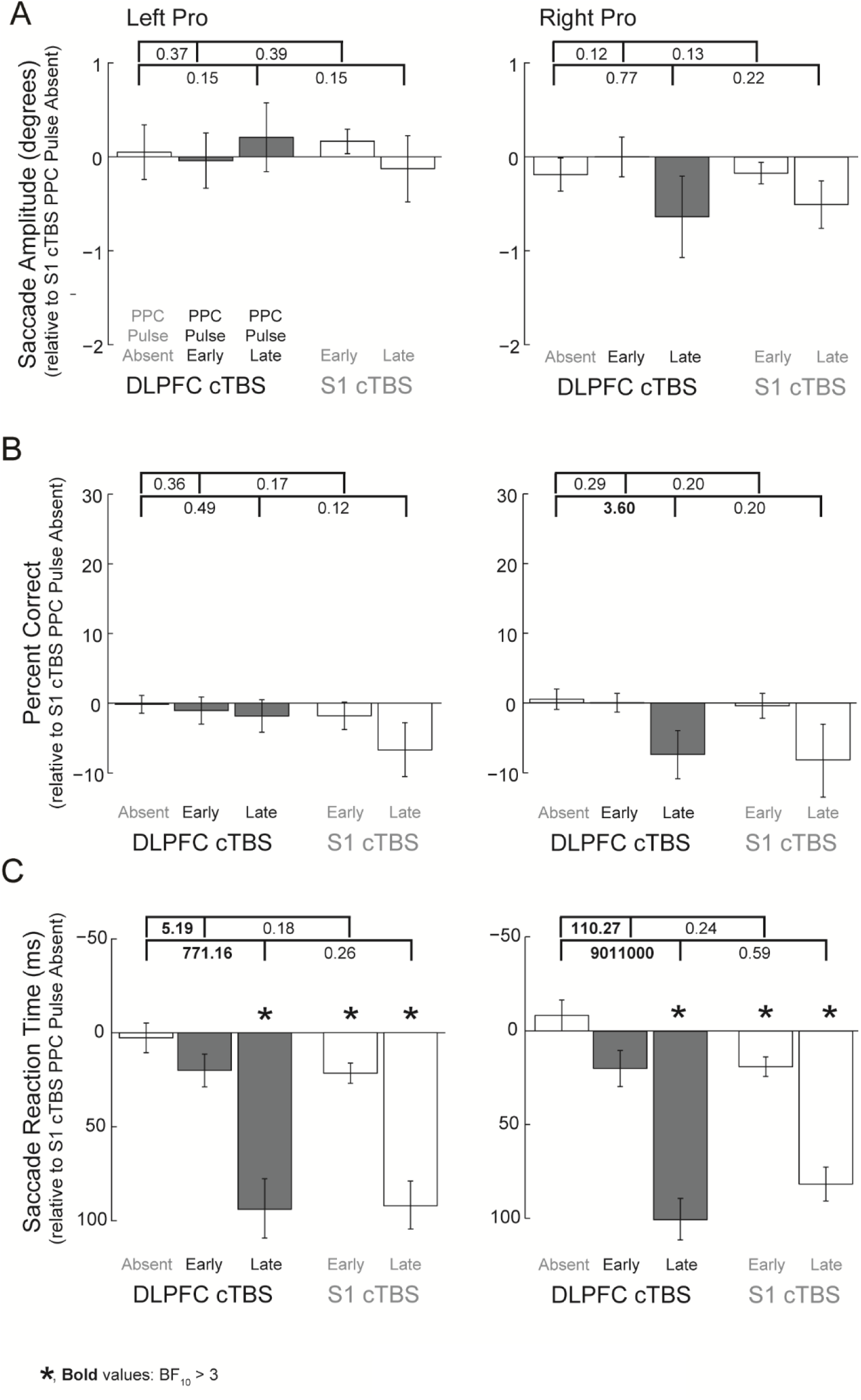
Effects on left and right pro-saccades when the double lesion involving DLPFC cTBS and PPC TMS is compared to the single lesion conditions. Conventions as in Figure 4.

#### Percentage correct direction

There was substantial evidence that the impairments to rightwards pro-saccades were greater following DLPFC cTBS when the late PPC pulse was present (Figure 7B; BF_10_ = 3.60), but no other evidence for impairments was substantial (Table 5B).

#### Saccade Reaction Times (SRT)

There was decisive evidence for reaction time impairments at the late PPC pulse time following DLPFC cTBS (Table 5C). Also, there was substantial evidence that the combined effects of DLPFC cTBS and PPC pulses resulted in enhanced impairments relative to DLPFC cTBS alone (Figure 7C).

#### Interpretation

As with FEF cTBS, DLPFC appears not to be critical to pro-saccade amplitudes. Interestingly, the late PPC pulse following DLPFC cTBS impaired rightward pro-saccade performance compared to DLPFC cTBS alone, but it is difficult to interpret this as compensatory as this condition did not actually produce substantial evidence for an impairment (BF10 < 3, Table 5).

## Conclusions

Our findings for a general lack of additive effects from two TMS lesions to critical nodes in anti-saccade programming suggest that these saccade behaviors are governed by distributed computations. A cTBS lesion to FEF, or DLPFC, may be interpreted as being consequential for the communication of information in a cortical network critical to mapping the spatial positions for an anti-saccades (Bullmore & Sporns, 2009; Sporns, Honey, & Kötter, 2007). The present study suggests DLPFC may be part of this network previously more emphasized to involve FEF and PPC (Medendorp et al., 2005; Moon et al., 2007; Munoz & Everling, 2004). Importantly, these three nodes have been shown to be part of interconnected frontoparietal networks which are recruited when attentional control is needed (Dosenbach, Fair, Cohen, Schlaggar, & Petersen, 2008; Ptak, 2012; Thiebaut de Schotten et al., 2011; Tschentscher, Mitchell, & Duncan, 2017; Vossel et al., 2014). Taken together, the evidence suggests that network interactions are important, over and above summated contributions of individual nodes.

